# An analysis of current state of the art software on nanopore metagenomic data

**DOI:** 10.1101/288969

**Authors:** Samantha C Pendleton

**Affiliations:** Institute of Mathematics, Physics and Computer Science, Aberystwyth University

**Keywords:** bioinformatics, nanopore, metagenomic

## Abstract

**Context:** Our insight into DNA is controlled through a process called sequencing. Until recently, it was only possible to sequence DNA into short strings called “reads”. Nanopore is a new sequencing technology to produce significantly longer reads. Using nanopore sequencing, a single molecule of DNA can be sequenced without the need for time consuming PCR amplification (polymerase chain reaction is a technique used in molecular biology to amplify a single copy or a few copies of a segment of DNA across several orders of magnitude).

**Aims:** Metagenomics is the study of genetic material recovered from environmental samples. A research team from IBERS (*Institute of Biological, Environmental & Rural Sciences*) at Aberystwyth University have sampled metagenomes from a coal mine in South Wales using the Nanopore MinION and given initial taxonomic (classification of organisms) summaries of the contents of the microbial community.

**Methods:** Using various new software aimed for metagenomic data, we are interested to discover how well current bioinformatics software works with the data-set. We will conduct analysis and research into how well these new state of the art software works with this new long read data and try out some recent new developments for such analysis.

**Results:** Most of the software we used worked very well: we gained understanding of the `ACGT` count and quality of the data. However some software for bioinformatics don’t seem to work with nanopore data. Furthermore, we can conclude that low quality nanopore data may actually be quite average.

## 1 Introduction

A wide variety of software is available today for bioinformatics. Majority of these tools that were created aim for short DNA reads, which researchers are familiar to working with.

Though new sequencing methods have made it possible to work with longer reads, making us curious if the bioinformatics tools also work with these data-sets. These significantly longer reads are going to be analysed with numerous bioinformatics tools; we want to know how the tools work and analyses nanopore data, specifically metagenomic data.

Using different software, we want to analyse the data to observe sequence similarities and find out what bacteria/species that resides within in mine. Our aim is to specifically observe quality, however we will also be looking into reads, `ACGT` content (in-depth `GC` analysis), plus time.

From a coal mine located in South Wales, DNA was extracted by a research team from the Institute of Biological, Environmental & Rural Sciences (IBERS); these mines had no internet access available.

The samples extracted were run in the nanopore technology, results were analysed in a pre-printed paper[4].

Two data sets were concluded from the mine expeditions; both from the same mine. There extracted 4 DNA samples for `BP_v1`, generated Dec-2016. 8 samples were extracted for `BP_v2`, generated Apr-2017. Version 2, `BP_v2`, is better quality as the researcher had optimised the DNA extraction to improve recovery yields from this kind of environment by using an improved protocol[3].

## 2 Method

A great deal of in-depth analysis of the data-sets will be conducted and discussed throughout. Whilst in the mine, the samples were run through the new MinION^1^ technology, developed by Oxford Nanopore^2^.

I downloaded source code of various bioinformatics tools and installed them, some were run through the use of the IBERS cluster server - looking into sequence similarity of the `FAST5`^3^ data, with the tools: `poretools`, `Goldilocks`, `FastQC` and `BLAST`.

### 2.1 poretools

`poretools`[7] is a toolkit for analysing nanopore sequence data - it was developed in Aug-2014 with 77 issues as of Sep-2017 on Github^4^ - possibly explaining potential errors and bugs.

#### 2.1.1 Read Lengths

`BP_v1` had more short reads, with a total of 1,770 files. Whilst `BP_v2` generated consistently longer reads and a total of 3,019 files - it was not quality filtered; it had MUX genes, which were later removed for analysis. In the histograms, we can see the comparison of the data-sets; they were recreated from the ones used in the IBERS pre-print paper[4].

**Figure 1:**
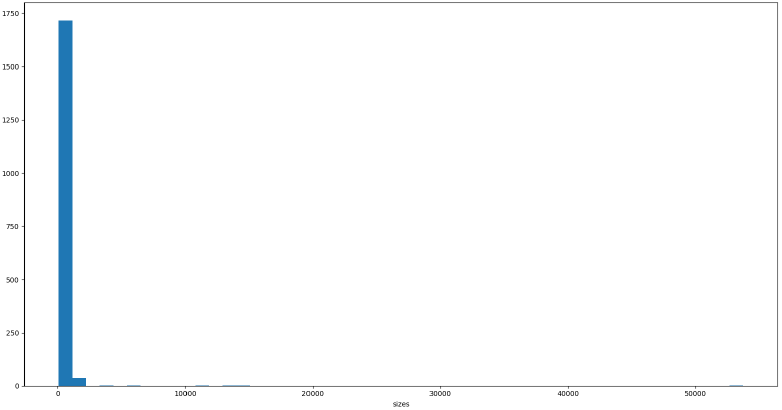
Histogram of `BP_v1`:- `x-axis`: read length (size); `y-axis`: cumulative frequency (count).

**Figure 2:**
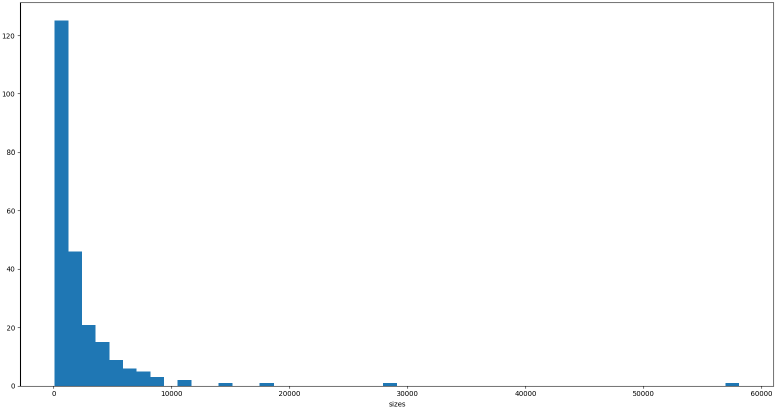
Histogram of `BP_v2`:- `x-axis`: read length (size); `y-axis`: cumulative frequency (count).

After some quality tests, it was observed that the longest reads were low in quality and so most likely errors whilst the samples were running through the MinION - we limited the data to 10,000 base pairs (bp) and compared the full data-set to the limited set frequently throughout.

**Figure 3:**
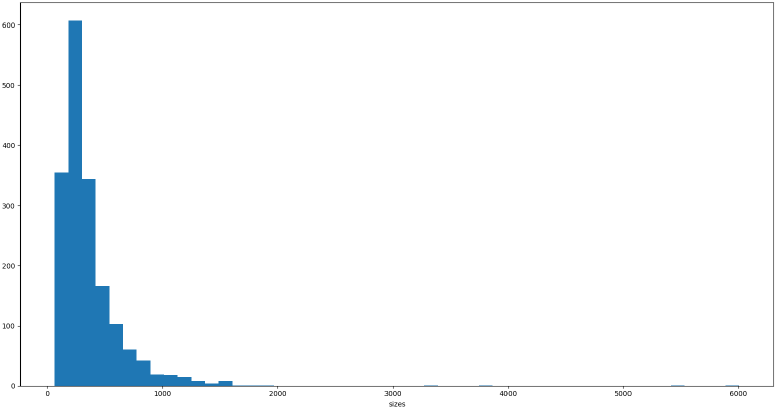
`BP_v1` limited to 10,000 bp:- `x-axis`: read length (size); `y-axis`: cumulative frequency (count).

**Figure 4:**
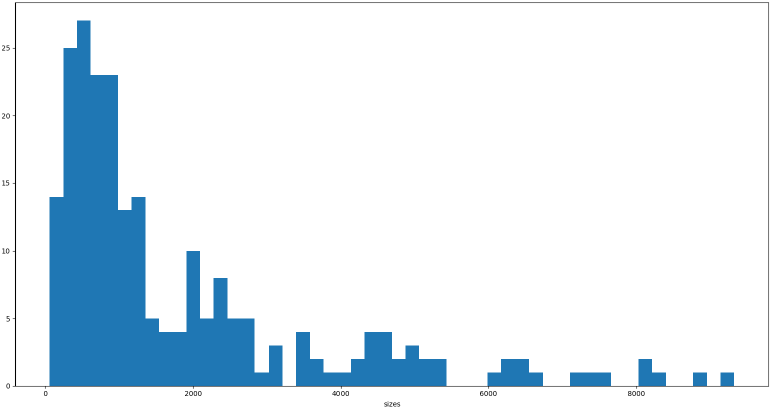
`BP_v2` limited to 10,000 bp:- `x-axis`: read length (size); y-axis: cumulative frequency (count).

#### 2.1.2 Quality

The quality of both data-sets averaged low with `BP_v2` being slightly higher in quality compared to `BP_v1`. `poretools`’ `qualpos` function produces box plots, these plots are useful displaying results of low quality, however, there’s an unusual number of plots scoring at 60: these are outliers which could be a result due to software not updated (as stated earlier the high number of issues on the Git repository).

**Figure 5:**
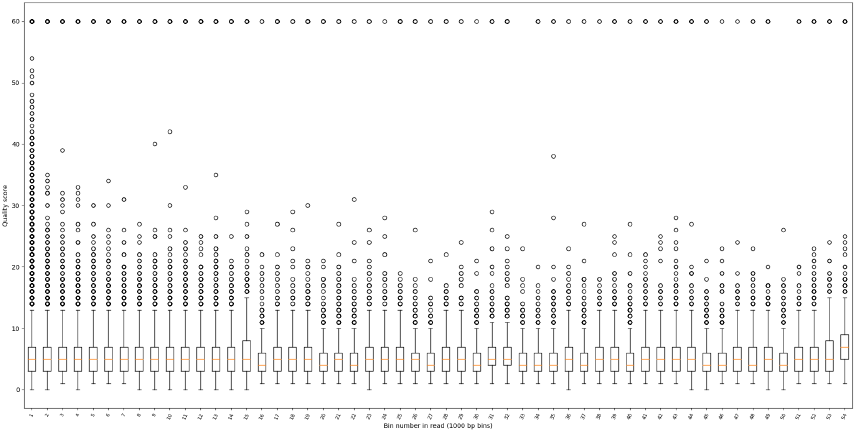
`poretools`’ use of `qualpos` of `BP_v1`:- `x-axis`: bin number in read; `y-axis`: quality score.

**Figure 6:**
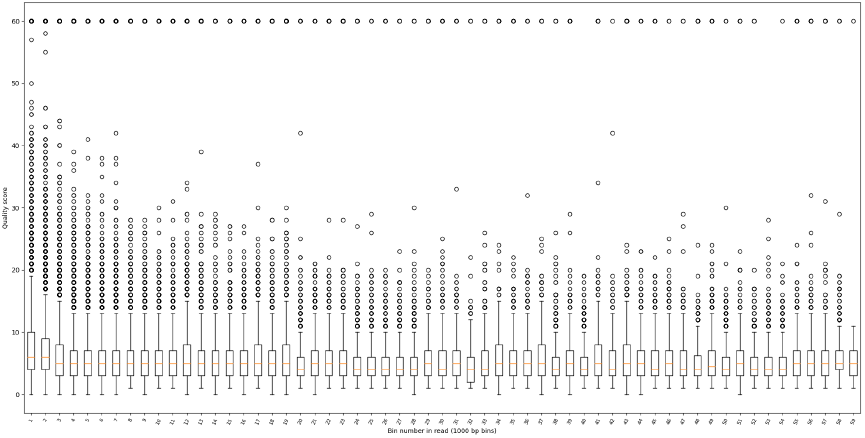
`poretools`’ use of qualpos of `BP_v2`:- `x-axis`: bin number in read; `y-axis`: quality score.

Using `poretools`’ `qualdist` (summary quality scores) function, we can observe the quality count: we can analyse the lack of count for letters: concluding further that both data-sets are poor.

##### Note

letters are what we are aiming for, these are good quality scores; ‘%’ is a specific symbol that relates to bad quality - both data sets are high in this symbol; count of 116,956 in `BP_v1` & 81,437 in `BP_v2`. We can see that `BP_v1` has a higher count of ‘bad’ symbols giving more evidence that `BP_v1` is lower quality than `BP_v2`.

**Figure 7:**
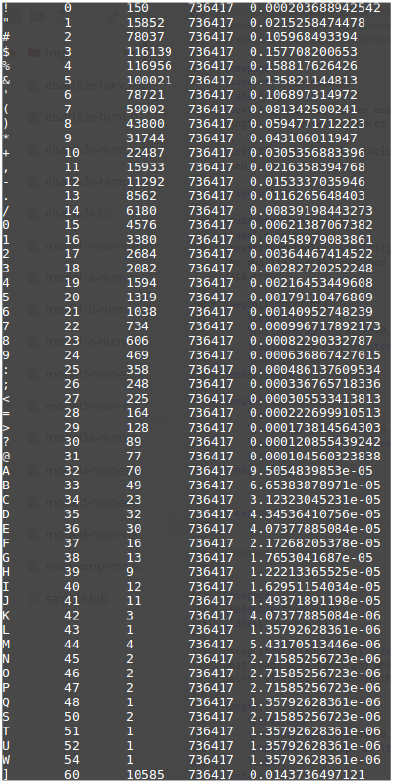
`poretools`’ qualdist of `BP_v1`.

**Figure 8:**
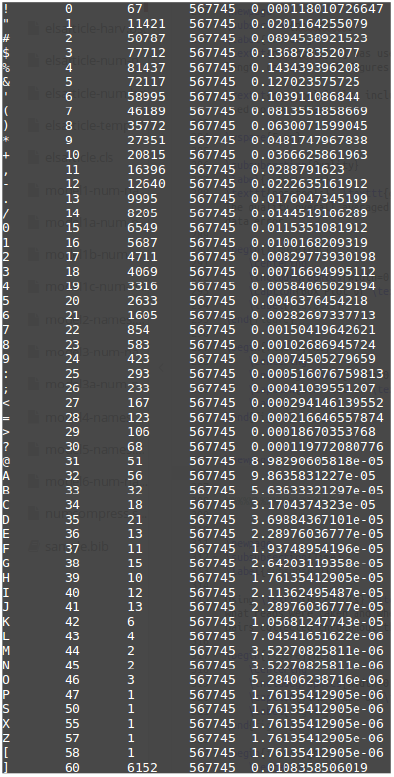
`poretools`’ qualdist of `BP_v2`.

#### 2.1.3 Time

The research team left the mine at 50 minutes - `BP_v1` was paused for 6 hours then continued at a researcher’s home. Meanwhile `BP_v2` ran throughout the return journey: DNA reads were affected after they left the mine during the elevator trip back to the surface plus the car journey (breaks, bumps, and going up/down a hill).

In the plots below, there are sections that show large vertical jumps, which are sudden production of base pairs: after studying, these are sections which were high in `AT` repetitive reads.

**Figure 9:**
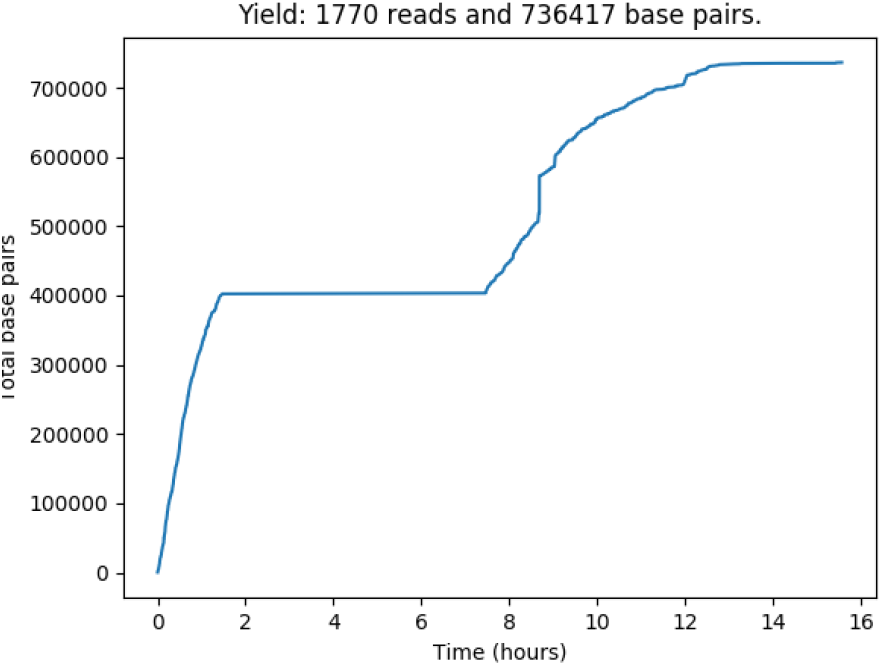
Time graph of `BP_v1` using `yield plot`:- `x-axis`: time (hours); `y-axis`: count of total base pairs.

**Figure 10:**
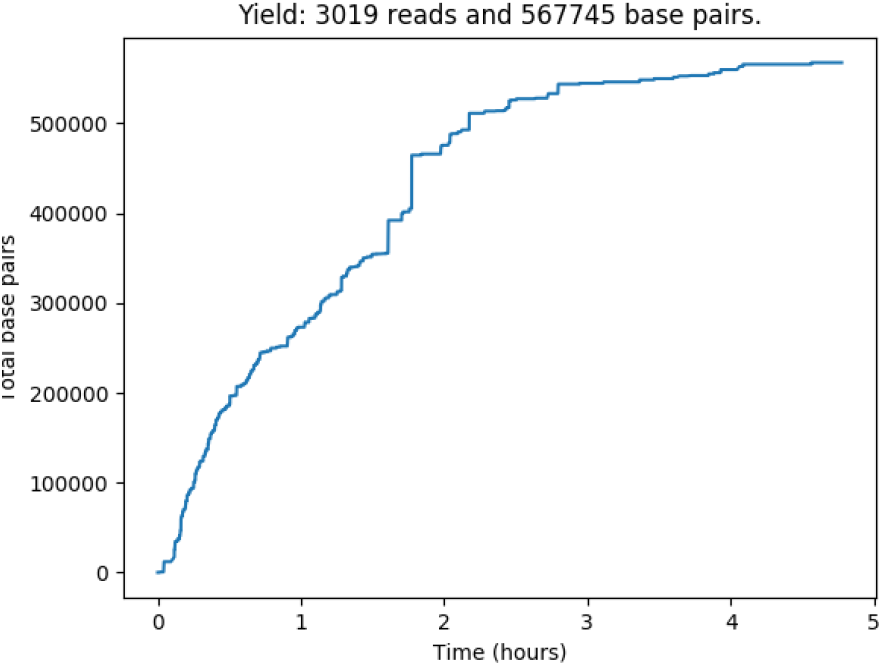
Time graph of `BP_v1` using `yield plot`:- `x-axis`: time (hours); `y-axis`: count of total base pairs.

There is an linear increase of base pairs as time increases for both data-sets, as we would expect (ignoring the pause for `BP_v1`). Both data-sets seem to have collected 400,000 base pairs within two hours - `BP_v1` collected 736,417 base pairs over 10 hours (16 hours total with a 6 hour pause) and `BP_v2` collected 567,745 in 5 hours.

We can see at 9 hours for `BP_v1`, there was a sudden long peak, reduced/no yield in DNA - this is potentially the longest read of the data set and coincidental A heavy.

The top 5 longest reads of `BP_v2` were taken after 50 minutes - we can bring facts together to conclude why there was data error: after leaving the mine, the team continued to run the analysis during the journey and and car journey affected the sequences.

However, other results from different tests told us how the top 5 T heavy reads of `BP_v2` were not the same as the top 5 longest reads. Moreover, the top 5 T heavy reads were majority after the first hour too, despite one read that was actually within the 50 minutes; after analysing this read through `BLAST`, we see that despite high in T, it actually had bacteria and fungus, but also sequence similarities of plant and animal.

### 2.2 Goldilocks

`Goldilocks`[9] was developed to “quickly locate *interesting* regions on the human genome that expressed a desired level of variability, which were ‘*just right*’ for later variant calling and comparison” - it was developed in Aug-2014 with last update Jul-2016 and 7 issues as of Sep-2017^5^.

The use of `Goldilocks` allowed us to observe `ACGT` content and `GC` in more detail through a scatter plot.

#### Note

`Goldilocks` produces linear graphs and the reads are in cardinal order: read as data position in file, which is most likely time.

#### 2.2.1 GC

GC in DNA analysis tells us about the stability: DNA with low `GC` content is less stable.

`A` *adenine* pairs with `T` *thymine*, and `C` *cytosine* pairs with `G` *guanine*. Majority of researchers look for higher ratio in `GC` compared to `AT`.

`BP_v1` is quite scattered with a cluster at location 100,000 - `BP_v2` average 0.7 `GC` content with an unusual section of low `GC` content at location 300,000.

**Figure 11:**
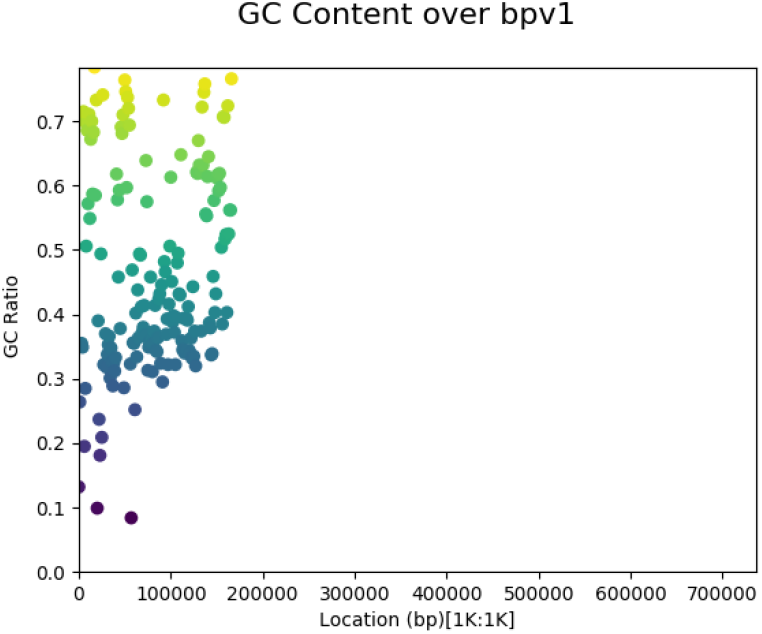
`GC` ratio of `BP_v1`:- `x-axis`: the location along the genome; `y-axis`: the value of the censused region.

**Figure 12:**
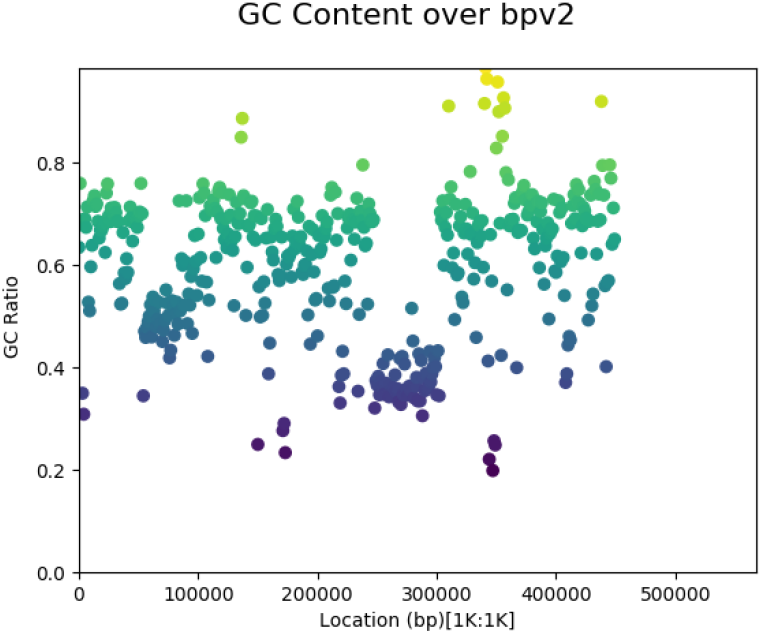
`GC` ratio of `BP_v2`:- `x-axis`: the location along the genome; `y-axis`: the value of the censused region.

#### 2.2.2 ACGT

The `NucleotideCounterStrategy` shows that both data-sets have a high number of `Ts` with `BP_v2` having a peak of `T` at base pair location 250000-300000; `BP_v1` has an unusual peak of `A` at location 75000-125000.

**Figure 13:**
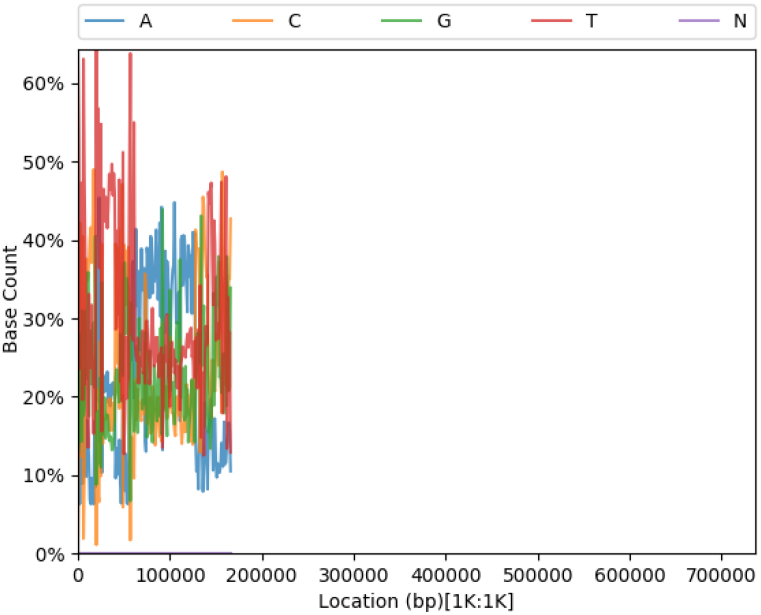
`ACGT` content of `BP_v1`:- `x-axis`: the location along the genome; `y-axis`: percentage of content cover.

**Figure 14:**
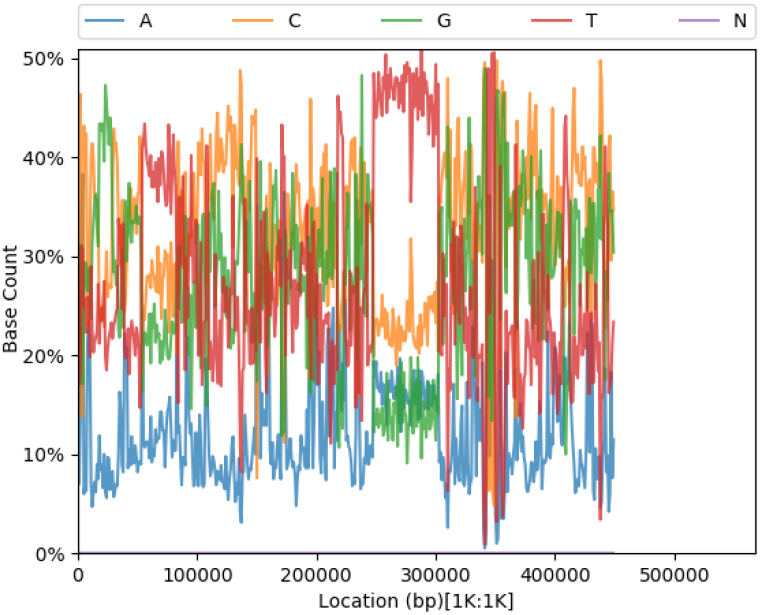
`ACGT` content of `BP_v2`:- `x-axis`: the location along the genome; `y-axis`: percentage of content cover.

`Goldilocks`’s `g.query` was used to find the reads with the most Ts in `BP_v2` & most `As` in `BP_v1` to which we then used the following command in a Linux terminal:

~~~
head -n10 BP_v2.fasta.fai
~~~

This method is to observe the top longest reads for both data sets, we did a quality check on both: the longest reads were poor in quality (to be explained further).

##### Note

to use the `FAST5` files with `Goldilocks`, we had to use `poretools` to convert them to `FASTA`^6^ - SAMtools[6] was used to index `FASTA` files so we could use them with `Goldilocks`

texttt`poretools` was also used to convert the `FAST5` files to `FASTQ`^7^ for use in `FastQC`.

### 2.3 FastQC

`FastQC`[1] was released Apr-2010, the oldest bioinformatics tool we are using, but it’s most recent updated was Mar-2016. It provides basic statistics of the datasets including read length and quality.

#### Note

`FastQC` has produced graphs that are not useful as the `x-axis` is uniform but stretched; whereas `Goldilocks` produces linear graphs: we want to observe read positions in linear form. Moreover, `FastQC` graphs do not display the whole data-sets; they are, unknown to us, limited.

#### 2.3.1 Read Length

Some statistical analysis of the data-sets that `FastQC` provides includes details of read length and `GC` content. There are 1761 total sequences in `BP_v1`, whilst `BP_v2` has 236 - `BP_v1` has more reads since they are shorter, `BP_v2` has less in total but they’re longer reads.

#### 2.3.2 GC

The `GC` ratio of `BP_v1` is 51%, whilst `BP_v2` is 60%. Comparing `GC` in `FastQC` & `Goldilocks` may be quite difficult here due to difference of graphs: `FastQC` plots are line whilst `Goldilocks` creates scatter.

**Figure 15:**
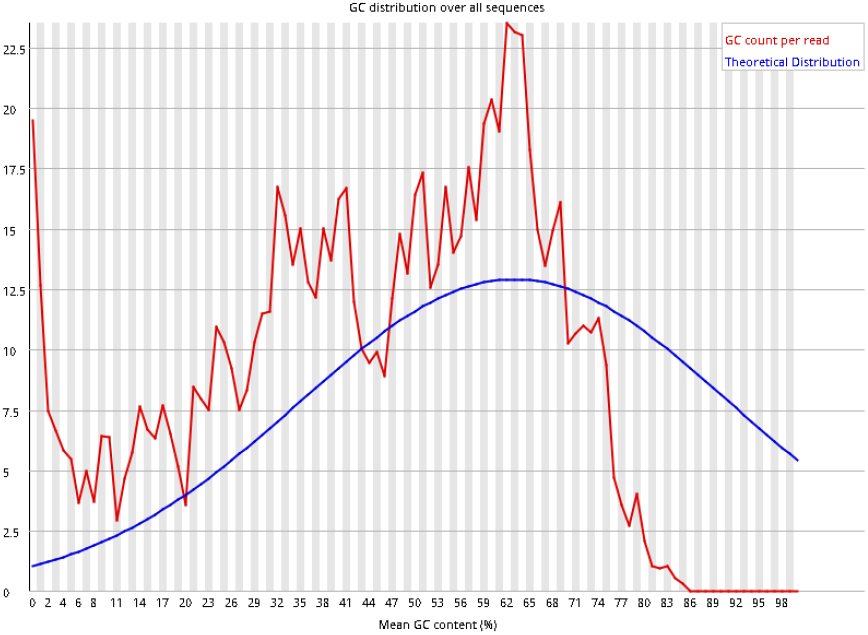
Plot of `GC` content of `BP_v1` in `FastQC`:- `x-axis`: mean content (%); `y-axis`: `GC` count per read.

**Figure 16:**
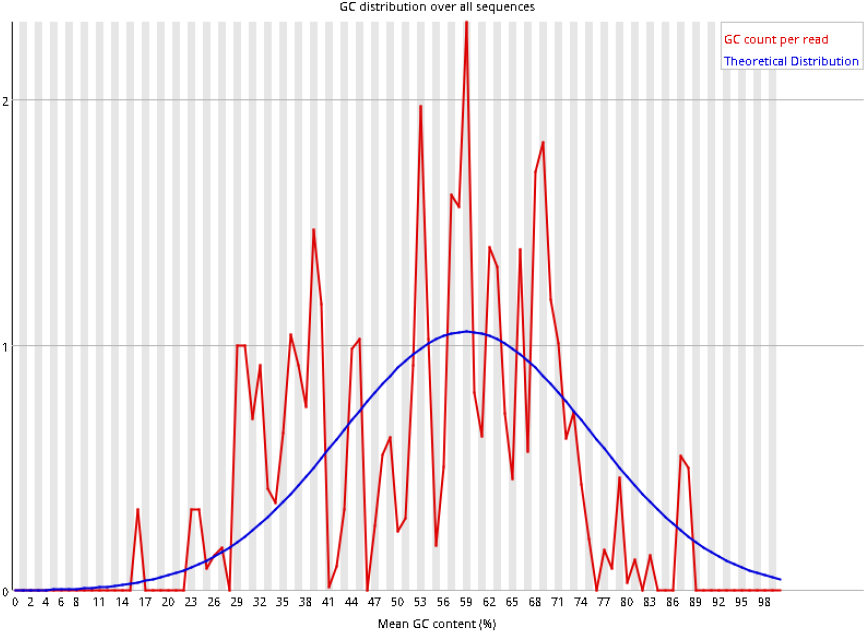
Plot of `GC` content of `BP_v2` in `FastQC`:- `x-axis`: mean content (%); `y-axis`: `GC` count per read.

#### 2.3.3 ACGT

Plotting `ACGT` content with `FastQC` produces truncated `x-axis`: they scale in an unusual way. Comparing to `Goldilocks`’ `ACGT` plots, we can see visible differences - T count in `BP_v2` steadily increases over time, as if logarithmic. These results of the `FastQC` plots are due to the truncated `x-axis` and it’s scaling.

**Figure 17:**
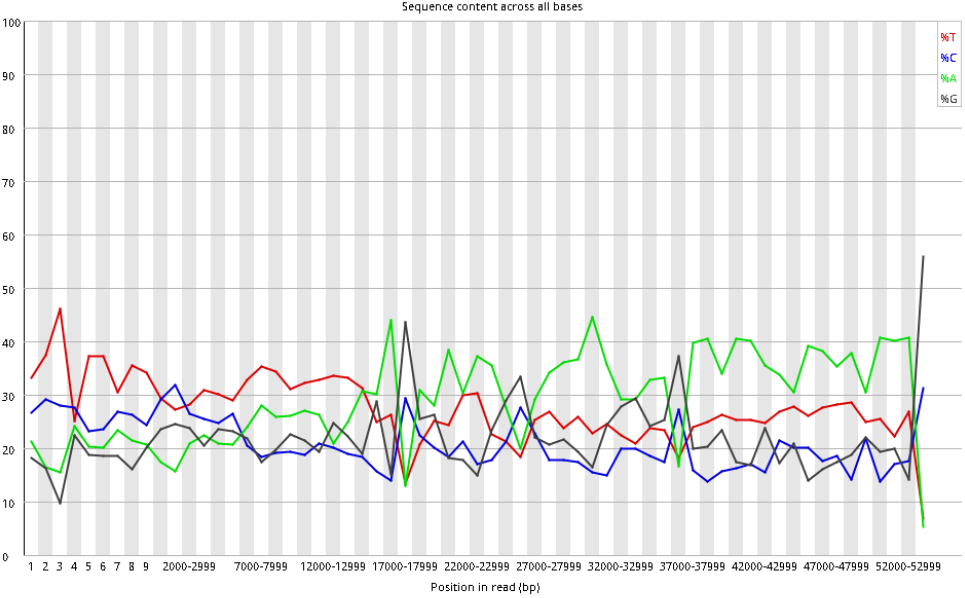
Plot of `ACGT` sequence content of `BP_v1` in `FastQC`:- `x-axis`: position in read (bp); `y-axis`: score.

**Figure 18:**
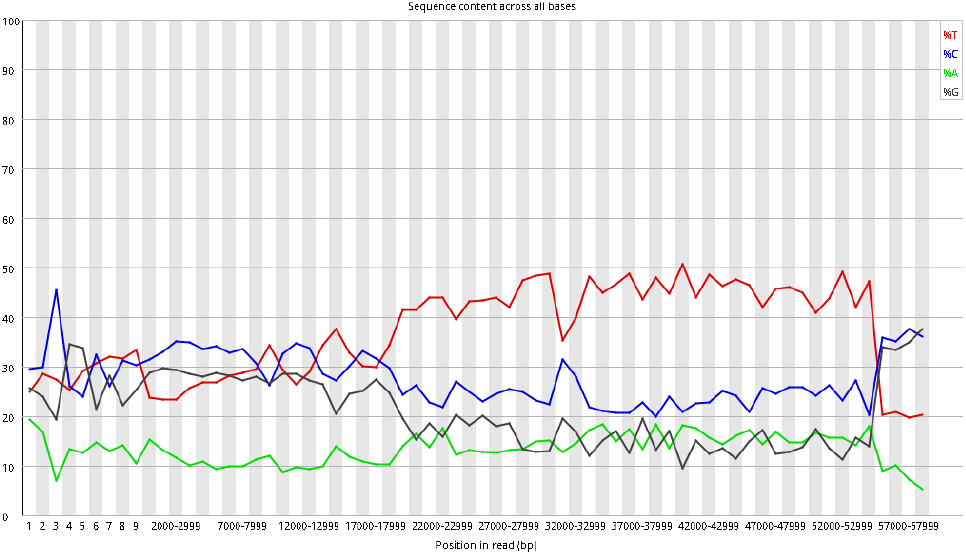
Plot of `ACGT` sequence content of `BP_v2` in `FastQC`:- `x-axis`: position in read (bp); `y-axis`: score.

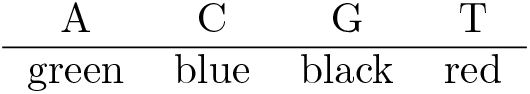

#### 2.3.4 Quality

Quality tests were conducted using `FastQC`, as we discovered earlier with `poretools` both `BP_v1` and `BP_v2` are low in quality. Despite the unusual `ACGT` plots of `FastQC`, the quality plots back-up the findings from `poretools`: despite `FastQC`’s results are limited from the `x-axis`, there are no outliers unlike `poretools`.

The test of quality are again displayed as box plots. The below plots are similar to `poretools`: low quality. However, when extracting some single reads of `BP_v2` and plotting, some values plotted in the medium range.

`FastQC` results are very useful, despite the limited box plots and strected x-axis. As stated earlier, `FastQC`’s x-axis is uniform but stretched, at first were misunderstood as logarithmic graphs; in these instances we prefer `Goldilocks`’ plots due to their graphs being linear.

The plots background colour scheme (observing data subsiding in the red area) is a unique feature.

**Figure 19:**
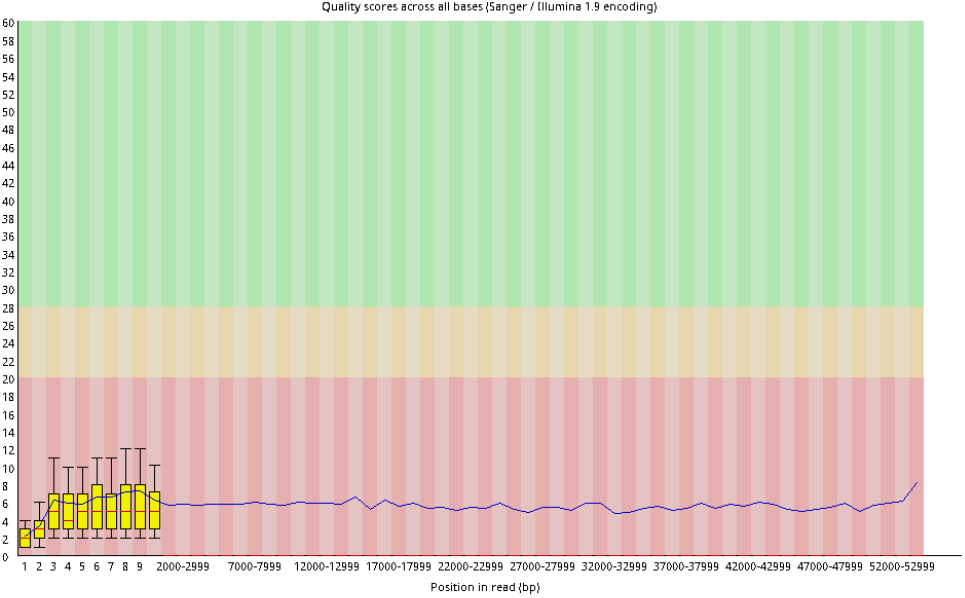
Plot of `BP_v1` reads in `FastQC` quality graph:-`x-axis`: position in read; `y-axis`: score.

**Figure 20:**
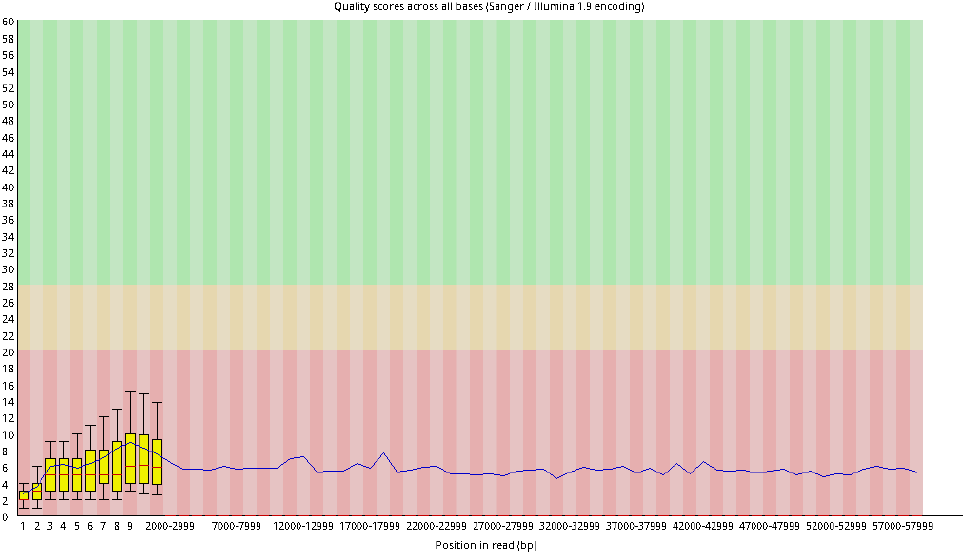
Plot of `BP_v2` reads in `FastQC` quality graph:-`x-axis`: position in read; `y-axis`: score.

As mentioned earlier, we had two versions of both data-sets - one original of both sets and the other were both sets limited to 10,000 base pairs.

When limiting the data-sets to 10,000 base pairs, we get the following plots below - `BP_v2`’s highest values on the box plots almost reach the medium/decent quality range - in a further study some random reads that were tested in `FastQC`’s quality test, we discovered that many ranged in the medium area - perhaps the data that is reliable was actually a medium score quality but majority of the data is low quality that it lowers the average.

**Figure 21:**
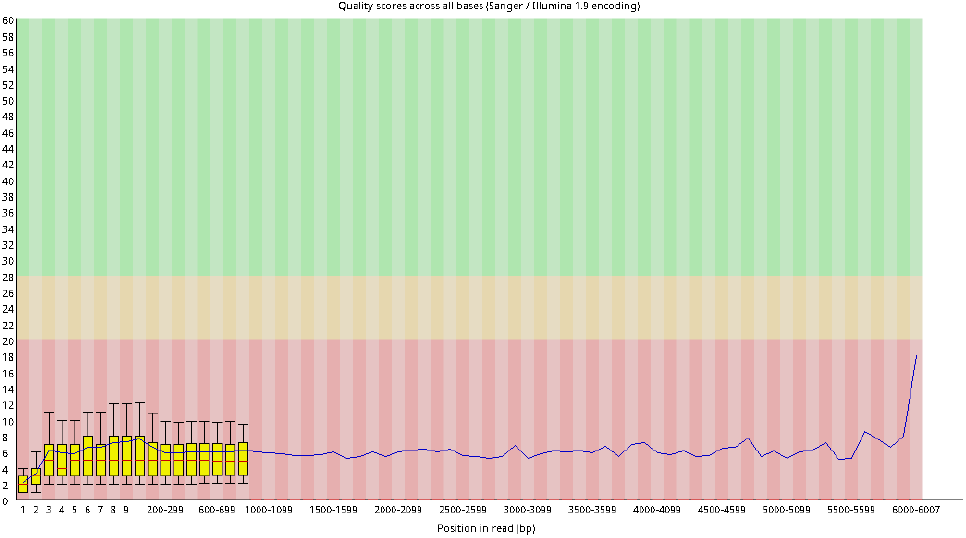
Plot of `BP_v1` reads in `FastQC` quality graph, limited to 10,000 bp:- `x-axis`: position in read; `y-axis`: score.

**Figure 22:**
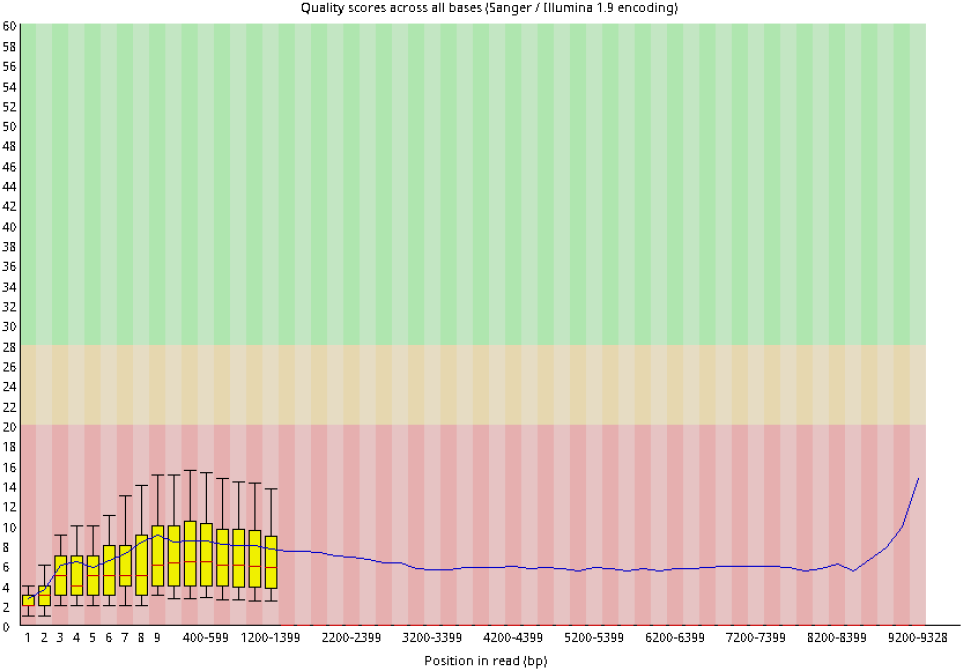
Plot of `BP_v2` reads in `FastQC` quality graph, limited to 10,000 bp:- `x-axis`: position in read; `y-axis`: score.

### 2.4 BLAST

`BLAST`^8^ for *Basic Local Alignment Search Tool* is an algorithm for comparing sequence information: the nucleotides of DNA sequences. We used this tool for analysing sequence similarities:- alignment lengths: aiming for results within hundreds; bit-score: high bitscores mean better sequence similarity; and percentage of identical matches. We used `BLASTn` on both data-sets and searched the results (taxon IDs) in the NCBI database.

#### Note

links in this section may have expired.

#### Longest reads

Further tests included using nucleotide `BLAST` (`BLASTn`) with `discontigious MEGABLAST` on the longest reads of both data sets to yet further prove the quality is low: the query cover was 0% of tiny hits, under 100 base pairs (bp).

~~~
BP_v1
~~~

Species: Japanese rice fish & Zebrafish (animals), Fruit flies (insects).

Dictyostelium discoideum: species of soil-living amoeba - commonly referred to as slime mould.

**Figure 23:**
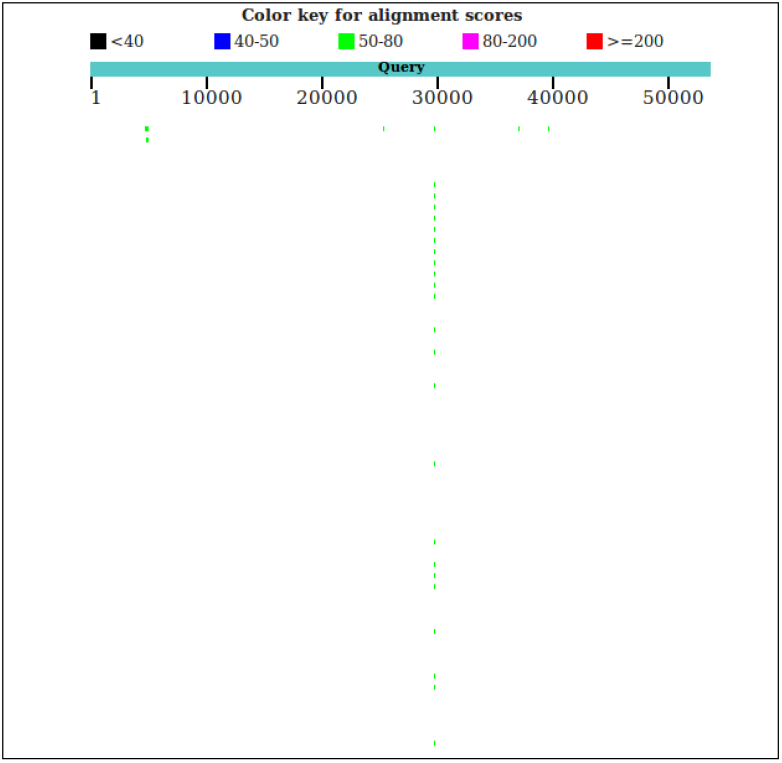
`BP_v1` `BLAST` alignment score of the longest read.

~~~
BP_v2
~~~

Species: Mouse & Tapeworm (animals), Tomato & Kiwi (plant), Fruit flies (insects), Human.

Parasites (including flatworms & malaria), Fungus

**Figure 24:**
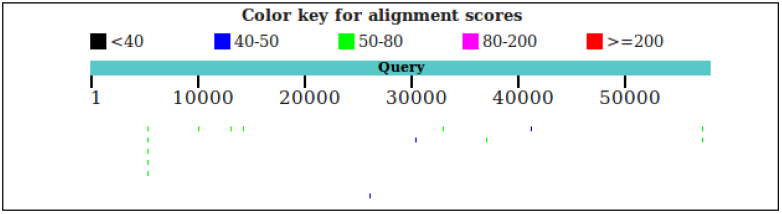
`BP_v2` `BLAST` alignment score of the longest read.

#### Random reads

Due to the long reads being low in quality and not many results of reliable species, further tests on the data sets were conducted: specifically we aim to `BLAST` random, shorter reads.

After extraction of random numerous shorter reads from `BP_v1` and the use of `BLAST`: no similarities appeared - the reads were mostly: `AT` content (short and repetitive).

Many random reads of `BP_v2` were tested and like results of `BP_v1`, no similarities appeared. However, after extracting a random read from `BP_v2` - we can observe this data read was bacteria, Anaerolinea thermophila^9^.

There was a query cover of 17%.

~~~
channel_349_bf5a3f39-3e35-4870-82eca7f9679d4000_qscore_9_read_score_-1.8
~~~

**Figure 25:**
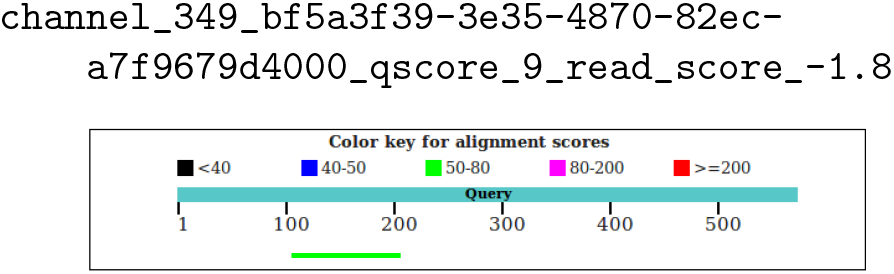
Alignment Score of a random `BP_v2` read.

**Figure 26:**
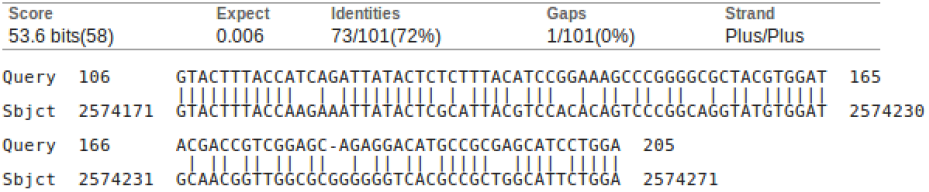
Alignment: Anaerolinea thermophila UNI-1 DNA, complete genome.

*Anaerolinea thermophilia* is thermophilic bacteria which thrives in high temperatures (41-122 degrees Celsius) - this result is quite unusual as the mine was not significantly at a high temperature; a researcher stated the mine was 15-20 degrees.

For further analysis, we extracted another random read from `BP_v2` (ID: 164). Despite no actually good hit, a reoccurring factor was E-coli.

The highest result of E-coli can be seen below: there was a query cover of 1% commonly.

~~~
channel_176_363eaef2-4772-408e-8641-5c1c111fd6bb_qscore_8.4_read_score_-1.6
~~~

**Figure 27:**
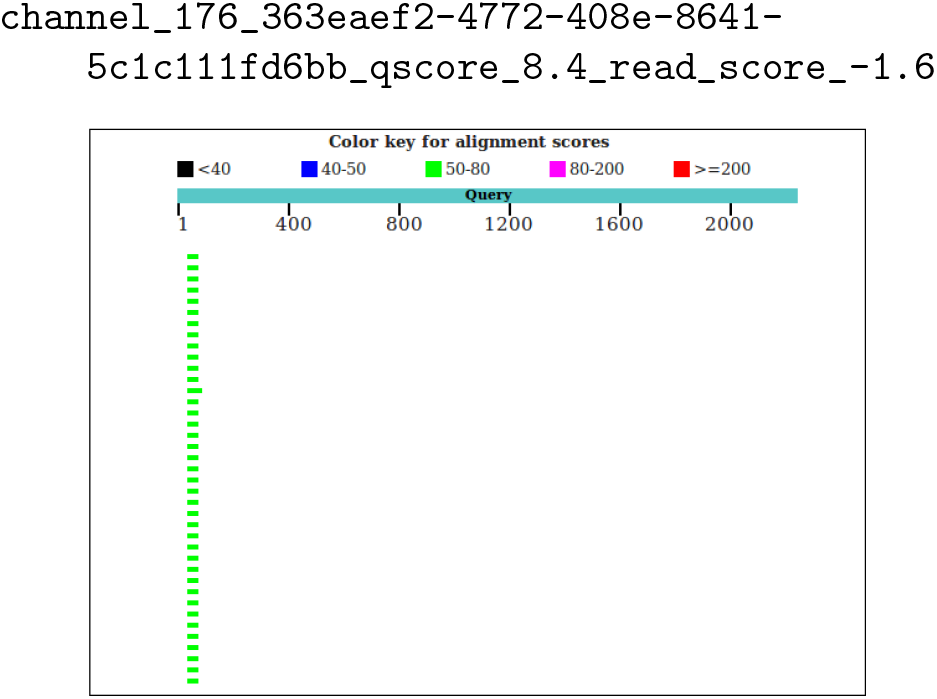
Alignment Score of a random `BP_v2` read.

**Figure 28:**
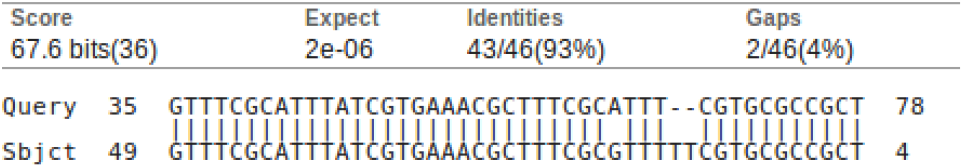
Alignment: Escherichia coli transposon Mu dl-R insertion site.

*Escherichia coli transposon*; a transposon is expected, it is a sequence of DNA that can move to new positions within the genome: *jumping genes*^10^. Furthermore, the alignment score seems to be 100: these are not genes however could be 10,000 matches of 1,000 genes.

#### Whole data-sets

Using AU IBERS cluster server, I was able to blast the data-sets as a whole, rather than just blasting long or random reads. We obtained a bunch of taxon IDs that we searched in NCBI database.

This method was used as the NCBI `BLAST` website would timeout due to the data-sets being large: CPU usage limit was exceeded. I had to `BLAST` locally with the `nt` nucleotide collection/database[2] (nr is the proteins collection).

`BP_v1` had poor results - unfortunately the highest result of alignment length was 49; highest bit-score was 67.6; and mostly 100% identical matches but this is because the reads are short.

#### Species found

⊲ Anaeromyxobacter dehalogenans 2CP-C: strain that efficiently reduces metals: ferric iron, Fe(III), and oxidised uranium (bacteria)
⊲ Capsicum annuum: Sweet and chili peppers (plant)
⊲ Pygocentrus nattereri: Red-bellied piranha (animal)

`BP_v2` had better results ranging from 300 to 900 in alignment score; bit-scores were as high as 440; and percentage of identical matches varied due to longer reads: for the top 5 highest alignment scores, the average result was 74.941%.

#### Bacteria found

⊲ *Neorhizobium galegae*: bacteria that forms nitrogen-fixing root nodules
⊲ *Rhodoplanes*: a phototrophic genus of bacteria - organisms that carry out photosynthesis
⊲ *Azorhizobium caulinodans*: bacteria that forms a nitrogen-fixing symbiosis with plants of the genus Sesbania (flowering plants in pea family)
⊲ *Bordetella flabilis*: recovered strains from cystic fibrosis (from human respiratory specimens)
⊲ *Nitrosomonas*: bacteria that oxidizes ammonia into nitrite as a metabolic process; found in areas that contains high levels of nitrogen compounds
⊲ *Rhodoplanes*: bacteria organisms that carry out photosynthesis
⊲ *Sideroxydans lithotrophicus*: bacterium isolated from iron contaminated groundwater

## 3 Discussion

The tools as a whole each had their own advantages; discussing `poretools`, as it is a software designed for nanopore data, it is handy with lots of features, the histograms plus other plots were very useful: we were able to observe the time plots, read length, and quality information with the data we gathered from the other tools. Though the outliers in the box plots don’t agree with the quality visualisations from `FastQC` so we aren’t able to detect whether this is due to `poretools`’ need for updating or something with `poretools` that we aren’t able to figure out.

`Goldilocks` on the other hand, was not designed for nanopore data however it works very well. If it were updated it could become a very practical tool for this new data. The use of `Python`^11^ made us able to alter the functions in our own way: `Matplotlib`^12^ enables us to alter the plots: making them log, changing colours, plus more - altering the `ACGT` plots with `Matplotlib` in `Python` allowed us the alter the colours of `GC` and observe the `AT` reads separately.

`FastQC` offers a variety, the basic statistical analysis it provides on the software is quite a handy observation. However, this bioinformatics tool is not suitable for nanopore data due to graph truncation. When shortening the reads to 10,000 bp, the plots seemed to be a little more relaible, however we still cannot understand why there are few box plots with a stretched `x-axis`.

`BLAST` is very functional and useful, but slow. `BLASTING` separate reeds is practical, however if the data is high quality the results would be much more reliable. As our data was low in quality, we had to `BLAST` random shorter reads, which came back with somewhat reliable results however we cannot continue blasting random reads out of the few thousands of reads in total.

Though despite `BLASTING` the data-sets as a whole taking quite some time, we were able to gain some good results, especially for `BP_v2`.

The more we see similarities in results from the different software, the more the can rely on them in future for nanopore data. When new bioinformatics tools are released, we can use the old tools as a comparison.

On the other hand, where there are differences, it is quite difficult to know whether which tool is more accurate. There are also external factors: if one tool has random outliers yet another does not, are these outliers correct or perhaps it’s a fault with the software? Plus we need consider the premises that if a software is faster than another, is it more reliable or precise?

## 4 Conclusions

I had aimed to demonstrate more software, however the `BLAST` jobs alone took many hours and another piece of software, `Centrifuge`[5], couldn’t run as it didn’t have enough RAM on the `IBERS` compute cluster to scale to the input data.

`Centrifuge` would have been a great bioinformatics tool to use as it is a tool for for metagenomic sequences^13^ - which is what we are working with.

Overall, the aim was to use these tools on longread data. Unfortunately the reads which were decent quality (plus good `BLAST` results) were short - so we couldn’t see what more the tools potentially had to offer for this data.

We can see that from the research, some of the tools work very well with this new nanopore data. We can use various features to make observations; coming to understand how nanopore quality is quite low, even in good data: however this may just be average quality score for nanopore data, some papers have Phred score of 10.53[8] - 1 in 10 error (see supplementary material: current version figure 1).

We realise that the minION is a very sensitive technology and running the DNA through the nanopore needs to be a steady procedure; otherwise results are affected greatly: the T count was very unusually high.

Many *state of the art* bioinformatics tools are not yet suitable for long-read data.

We take away from this research with an idea of bioinformatics tools and how well they work with this new long-read sequence data. Perhaps we can acknowledge the early state of nanopore sequencing itself - it is still biologically difficult to handle DNA and keep it intact for sequencing.

## Acknowledgements

I would like to thank Amanda Clare, Senior Lecturer at the Institute of IMPACS (*Institute of Mathematics, Physics and Computer Science)*, my supervisor throughout this project for the constant guidance and support.

I would also like to acknowledge, Andre Soares, PhD student in Geomicrobiology at IBERS (*Institute of Biological, Environmental & Rural Sciences*), who was a research member a part of the team who went in the mine - he assisted me throughout and provided information about the datasets whenever I asked.

Finally, I want to express appreciation for Samuel M Nicholls, PhD student of Bioinformatics at IBERS and the Department of Computer Science, who creator of `Goldilocks` - he helped with installation of some tools.

## Footnotes

1 MinION, “the only portable real-time device for DNA and RNA sequencing.” https://nanoporetech.com/products/minion

2 Oxford Nanopore, “enable the analysis of any living thing, by any person, in any environment”. https://nanoporetech.com

3 `FAST5`, “file-type in bioinformatics from from Oxford Nanopore”. http://bioinformatics.cvr.ac.uk/blog/exploring-the-fast5-format/

4 `poretools`, “a toolkit for working with nanopore sequencing data from Oxford Nanopore”. https://github.com/arq5x/poretools

5 `Goldilocks`, “locating genetic regions that are just right”. https://github.com/SamStudio8/goldilocks

6 `FASTA`, “file-type in bioinformatics”. https://zhanglab.ccmb.med.umich.edu/FASTA/

7 `FASTQ`, “file-type storing the Phred qualities”. http://maq.sourceforge.net/fastq.shtml

8 `BLAST`, “finding regions of similarity between biological sequences”. https://blast.ncbi.nlm.nih.gov/Blast.cgi

9 *Anaerolinea thermophila*, `NCBI` description of filamentous thermophilic bacteria. https://www.ncbi.nlm.nih.gov/pubmed/14657113

10 *Transposons*, “The Jumping Genes”. https://www.nature.com/scitable/topicpage/transposons-the-jumping-genes-518

11 `Python`, Programming Language. https://www.python.org

12 `Matplotlib`, `Python` 2D plotting library. https://matplotlib.org

13 `Centrifuge`, “Classifier for metagenomic sequences”. https://github.com/infphilo/centrifuge

